# Sub-membrane actin rings compartmentalize the plasma membrane

**DOI:** 10.1101/2023.08.22.554239

**Authors:** Jakob Rentsch, Selle Bandstra, Batuhan Sezen, Philipp Stephan Sigrist, Francesca Bottanelli, Bettina Schmerl, Sarah Shoichet, Frank Noé, Mohsen Sadeghi, Helge Ewers

## Abstract

The compartmentalization of the plasma membrane is a fundamental feature of cells. The diffusivity of membrane proteins in the plane of the membrane is significantly lower in cells than observed in artificial membranes. This seems due to the sub-membranous actin cortex, but tractable assays to prove a direct dependence of membrane protein motion on actin filaments have been elusive. We recently showed that a periodic array of sub-membrane actin filaments that surround neuronal axons in ring-like structures in the axonal initial segment (AIS) confines membrane protein motion between them, but the local enrichment of ion channels may offer an alternative explanation for compartmentalization. Here we show using computational modeling that in contrast to actin rings, the dense array of ion channels in the AIS cannot mediate membrane protein confinement. We go on to show that indeed the actin rings are closely apposed to the plasma membrane, and that they confine membrane protein motion between them in a ∼ 200 nm periodic pattern in progenitor-derived neuronal cells. We find that this compartmentalization is also detectable for inner-leaflet membrane proteins and multi-spanning receptors. Strikingly, several cell types with actin rings in their protrusions, like progenitor-derived astrocytes and oligodendrocytes, also exhibit lateral confinement of membrane proteins in a periodic pattern consistent with actin ring spacing. Actin ring-mediated membrane compartmentalization is thus not unique to neurons. Finally, we show that acute actin disruption in live progenitor-derived neuronal cells leads to a loss of membrane compartmentalization in areas that were compartmentalized before treatment. Taken together, we here develop a much-needed tractable experimental system for the investigation of membrane compartmentalization and show that actin rings compartmentalize the plasma membrane of cells.

## Introduction

The plasma membrane is according to the Singer-Nicolson fluid mosaic model (Singer and Nicolson, 1972) a continuous membrane bilayer in which membrane proteins are suspended. Saffman and Delbrück (Saffman and Delbrück, 1975) offered a quantitative model to describe the two-dimensional free diffusion of membrane proteins in fluid membranes, which depends on the viscosity of the membrane and the surrounding solvent as well as the geometry of the protein and the membrane, and very well describes the motion of transmembrane proteins in artificial bilayers (Weiß et al., 2013). However, in cellular membranes, membrane proteins mostly do not seem to undergo unhindered lateral diffusion. Rather, they exhibit diffusion coefficients about an order of magnitude lower than measured in artificial membranes (Kusumi et al., 2005). Based on such observations, it has been proposed that membrane proteins are confined into membrane subdomains formed by the cytoskeleton (Kusumi et al., 2005). Here, transmembrane proteins may be corralled by a mesh of sub-membrane actin filaments that act as physical obstacles to the motion of their intracellular portion and form small domains according to the “fence” model of membrane partitioning (Kusumi et al., 2005). On the other hand, lipids and lipid-anchored proteins, which are also found to exhibit subdiffusive motion cannot be confined in this way. However, their confinement may be explained by an array of transmembrane proteins anchored along the actin filaments in the meshwork that by occupying a fraction of membrane space act as obstacles and are sufficient to confine molecules to compartments in the “picket” model (Kusumi et al., 2005).

However, due to the dynamic nature of the sub-membrane actin meshwork, no tractable model for the mechanistic investigation of the “fence” nor “picket” model are available. This problem may be overcome after the discovery of a periodic network of actin rings spaced every 200 nm along neuronal axons (Xu et al., 2012). In contrast to cortical actin, which is highly dynamic (Gowrishankar et al., 2012), these rings are stable in their location (Albrecht et al., 2016) and exist for longer periods of time even in the presence of some actin-depolymerizing drugs (Leterrier et al., 2015). Indeed, we could previously show, that the diffusion coefficient of a lipid-anchored molecule is abruptly reduced over the time course of establishment of these actin rings in the axon initial segment (AIS) of developing neurons (Albrecht et al., 2016). Strikingly, in areas of reduced diffusion coefficient, correlative high-density single particle tracking (SPT) of membrane proteins and super-resolution microscopy of actin revealed that membrane protein motion was confined between actin rings in the AIS (Albrecht et al., 2016). However, since the AIS is the site of a strong developmental accumulation of ion channels between the actin rings (Brachet et al., 2010; Garrido et al., 2003; Zhang et al., 1998; Hedstrom et al., 2008; Gasser et al., 2012), it remains uncertain, whether the membrane compartmentalization is due to the actin rings or to the high local density of transmembrane ion channels (Huang and Rasband, 2016; Leterrier, 2018). Importantly, since then, actin rings have been reported over a large variety of cell types across the neuronal lineage (Hauser et al., 2018; He et al., 2016; D’Este et al., 2015, 2016; Zhong et al., 2014). They may thus present a tractable experimental system for the investigation of membrane compartmentalization by the cytoskeleton. The answer to this open question is fundamental to our understanding of membrane compartmentalization in neurons and in plasma membranes in general.

Here we ask if the sub-membrane actin rings alone can partition the plasma membrane using computational modeling and high-density, high-speed SPT in a variety of cell types. We find that wherever in the neuronal lineage we observe periodic actin rings, these compartmentalize the motion of membrane proteins. STED microscopy in live cells shows that actin rings are closely apposed to the plasma membrane. Using molecular inhibitors, we find that acute actin disruption in live cells abolishes membrane compartmentalization. Taken together our results establish actin rings as causative to membrane compartmentalization in a variety of cell types, suggesting a global mechanism for regulation of membrane protein motion in cells.

## Results

We first aimed to ask, whether the observed membrane partitioning in the AIS was due to actin rings or the local density of ion channels. To do so, we performed 3D high-speed, high-density SPT of GPI-GFP in DIV7 rat hippocampal neurons using anti-GFP nanobody (NB) conjugated quantum dots (QDs). When we plotted all individual detection events, they clearly accumulated in a periodic 200 nm spaced pattern, resembling stripes perpendicular to the direction of propagation of the axon (Figure 1 b-e) similar to what we previously observed (Albrecht et al., 2016). These stripes line the perimeter of the axon and appear especially striking when observed in a 3D rendering of the GPI-GFP localizations (Movie S1). We hypothesize that the observed pattern can be explained by either of two possible mechanisms: (i) the periodic actin rings could cause membrane compartmentalization, and (ii) the enrichment of large ion channels at the AIS may trap and accumulate membrane proteins between actin rings (Figure 1 f). We investigated three models for the location of the diffusion barrier: actin rings (Figure 1 g), a sparse (Figure 1 h) and a dense accumulation of channels proteins (Figure 1 i).

**Figure 1.**
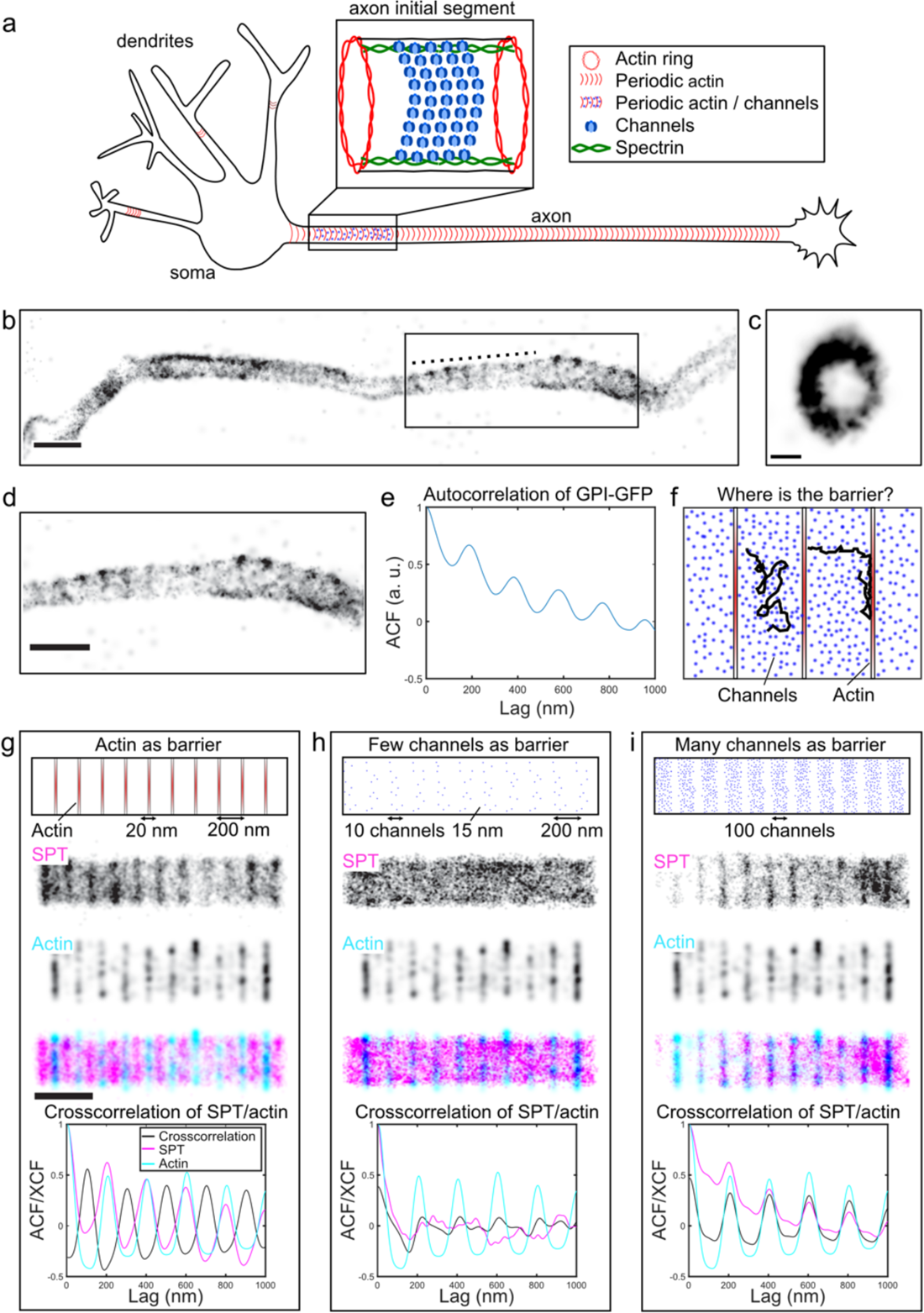
Simulations of SPT experiments reveal that actin rings act as a diffusion barrier. **(a)** Schematic representation of the cytoskeleton in neurons: the axon initial segment contains periodically organized actin rings (∼200 nm), which are interspersed by accumulations of channel proteins **(b)** GPI-GFP forms membrane domains in DIV7 rat hippocampal neurons. Reconstruction of localizations of SPT experiment of GPI-GFP tagged with QD’s (5000 frames, 200 Hz) **(c)** Projection along the dotted line in (b). Scale bar is 100 nm. **(d)** Zoom in of box shown in (b) **(e)** Autocorrelation along the neuronal process shown in (c) reveals periodicity of GPI-GFP domains at ∼200 nm. **(f)** Schematic representation of the two competing hypotheses: Actin acts as a diffusion barrier or channel proteins act as a diffusion barrier. **(g)** Simulations of SPT experiments in Fluosim. **(top)** Model geometry mimicking the organization of actin rings. **(middle top)** reconstruction of localizations of *in silico* SPT experiments over the geometry. **(middle)** *in silico* STORM reconstruction of actin rings. **(middle bottom)** Merge of SPT reconstruction and STORM of actin **(bottom)** Crosscorrelation of SPT of GPI-GFP and STORM of actin reveals that GPI-GFP forms domains located between actin rings. **(h)** Same as (f) but with few channel proteins as a model geometry: No accumulation of GPI-GFP into membrane domains can be observed. **(i)** same as (f) but with many channel proteins as model geometry: Accumulations of GPI-GFP into membrane domains can be observed but are localized on top of actin rings (in contrast to experimental data). If not otherwise indicated scale bars are 500 nm.

To address which of these models best explains our experimental observations, we performed simulations of single particle motion using the simulation software Fluosim (Lagardère et al., 2020). We used this software to generate synthetic time-series of images of single molecule peaks using parameters as found in cell experiments. We then added noise and motion blur to the level measured in real experiments and used single molecule localization microscopy software to analyze the resulting image stacks. We found that the synthetic data strongly resembled real measurements (Movie S2, S3), giving us confidence that this system would allow us to ask specific questions concerning compartmentalization of membrane molecule motion by obstacles. We went on to investigate three simulated scenarios mimicking the developmental time course of axon initial segment establishment (Figure 1a), where first actin rings appear, and subsequently ion channels become highly enriched over the following days. We thus first created a model geometry consisting of 20 nm thick lines that created 200 nm sized compartments between them to represent the actin rings (Xu et al., 2012). When we then generated artificial SPT movies of molecules diffusing over the model geometry, we found compartmentalization between the “actin rings” as repetitive peaks in an autocorrelation function beginning at a 70% probability for molecules to cross the “actin rings” (Figure S1). This is consistent with previous work suggesting that membranes can be compartmentalized even by incomplete “palisades” of transmembrane proteins anchored to sub-membrane actin filaments (Fujiwara et al., 2002). Furthermore, we simulated single molecule localization microscopy (SMLM) reconstructions of the “actin rings” and correlated the resulting pattern with the reconstructions of the simulated SPT data from the same simulation (Figure 1g). In this first simulation, we observed a clear pattern of SPT localizations, spaced between adjacent actin staining like we found in our experimental data before (Compare Figure 1b-e and Figure 1g and (Albrecht et al., 2016). Secondly, we generated model geometries consisting of a randomly spaced accumulation of circular exclusions zones with a diameter of 15 nm located between the periodic actin rings, the exclusion zones here mimicking large ion-channels anchored at spectrin/ankyrin between actin rings in the AIS. These exclusion zones reject molecules in 100% of encounters, but in this case, “actin” was 100% permeable. To simulate the increasing enrichment in the AIS over time (Jones et al., 2014), we performed simulations with 10 and 100 “ion channels” confined in this manner. When we then performed simulated SPT experiments of membrane molecules and SMLM experiments of actin as described above for these conditions, we could not detect an accumulation of membrane protein localizations between “actin rings” (Figure 1h, i). Strikingly in the scenario with 100 “ion channels”, molecules rather became accumulated on simulated actin staining (Figure 1i). Taken together, our simulations show that even very dense arrays of transmembrane proteins will not trap transmembrane proteins, but rather exclude them. Our simulations indeed support a model where the plasma membrane is compartmentalized by sub-membrane actin rings.

We next aimed to determine, whether the actin rings were indeed located in close proximity to the plasma membrane (Figure 2d). To do so, we performed line-scans across the equator of processes in live progenitor-derived neuronal cells labelled with SiR-actin and GPI-GFP (Figure 2a-c). We imaged SiR-actin in STED and GPI-GFP in confocal microscopy. When we quantified the distance between the peaks representing the GPI-GFP in the bilayer and the actin rings (SiR-actin) in many cells, we found that they were on average separated by around 30 nm. At the same time, when we measured the distance between membrane stainings by GPI-GFP and cholesterol-Abberior Star Red, we found a distance of around 11 nm. The localizations of multicolor fluorescent beads measured in the same assay yielded an accuracy of ∼ 2 nm (Figure 2e). This suggests that the actin rings are indeed very close to the plasma membrane, possibly close enough to anchor transmembrane proteins as “pickets” that act as obstacles to lateral membrane molecule motion.

**Figure 2:**
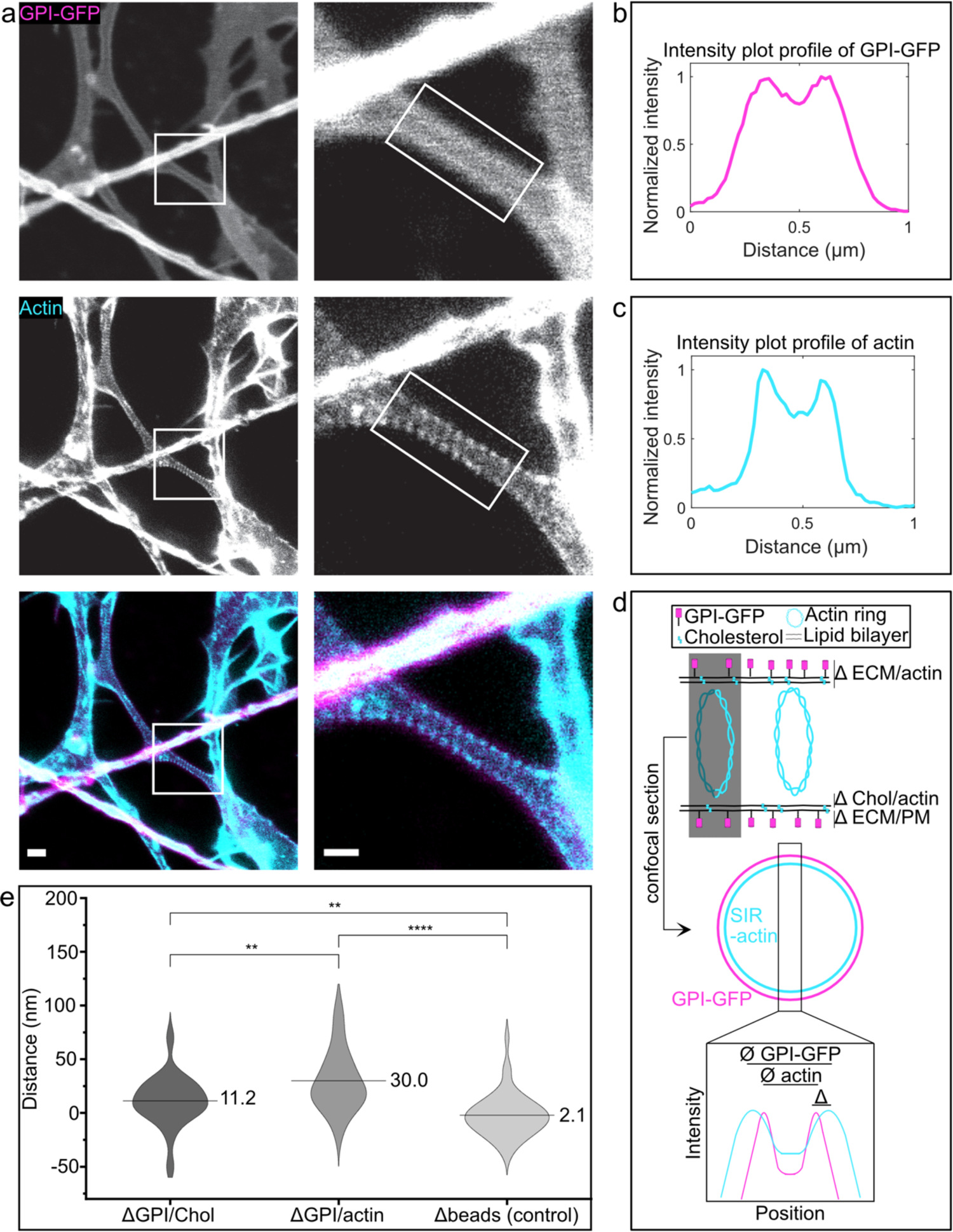
Actin rings are close to the plasma membrane. **(a) (top)** Confocal image of GPI-GFP **(middle)** Live-STED image of actin rings in progenitor-derived neuronal cells **(bottom)** Merge of actin and GPI-GFP. Right images are zoom ins of box on the left. Scale bars on the left are 1 µm. Scale bars on the right are 500 nm **(b)** Line profile of box of confocal image of GPI-GFP shown in (a) on the right **(c)** Line profile of box of live-STED image of actin rings shown in (a) on the right **(d)** Schematic representation of the calculation of the distance between GPI-GFP and actin rings: The distance of the peaks shown in (b) and (c) is measured. The difference between the distances is used to calculate the distance between actin rings and GPI-GFP **(e)** Violin Plot of the measured distances between GPI-GFP and the plasma membrane stained via Cholesterol-Star Red, GPI-GFP and actin rings and a bead control.

Next, we aimed to ask, whether an AIS is required for membrane compartmentalization. To do so, we used adult hippocampal neuronal progenitor cells (AHNPCs). They can be induced to differentiate into different types of cells of the neuronal lineage that all exhibit periodic actin rings (Hauser et al., 2018). Indeed, when we differentiated AHNPCs as described (Peltier et al., 2010), neuronal cells did not show an AnkG accumulation characteristic for an intact axon initial segment after DIV7 nor DIV14 (Figure S2). Since AnkG is required to target voltage-gated channels to the AIS (Garrido et al., 2003; Leterrier et al., 2017; Gasser et al., 2012) positioned by spectrin C-termini between the actin rings (Xu et al., 2012) we conclude here that neuronal cells derived from AHNPCs do not contain voltage-gated ion channel proteins in the characteristic dense organization present in the AIS. If membrane protein motion is compartmentalized between the periodic actin rings in these cells, it can thus not be controlled by an ion channel density. When we tracked GPI-GFP diffusion via QDs in these cells (Figure 3a), we still found that they formed repetitive ∼ 200 nm size domains. (Figure 3b,c) Strikingly, the observed membrane domains seemed to be stable over time as the same areas were visited multiple times during the observation period (Figure 3e). GPI-GFP in membrane domains remained mobile (D = 0.04 µm^2^s^-1^) and exhibited only slightly subdiffusive motion (α = 0.9, Figure S3).

**Figure 3:**
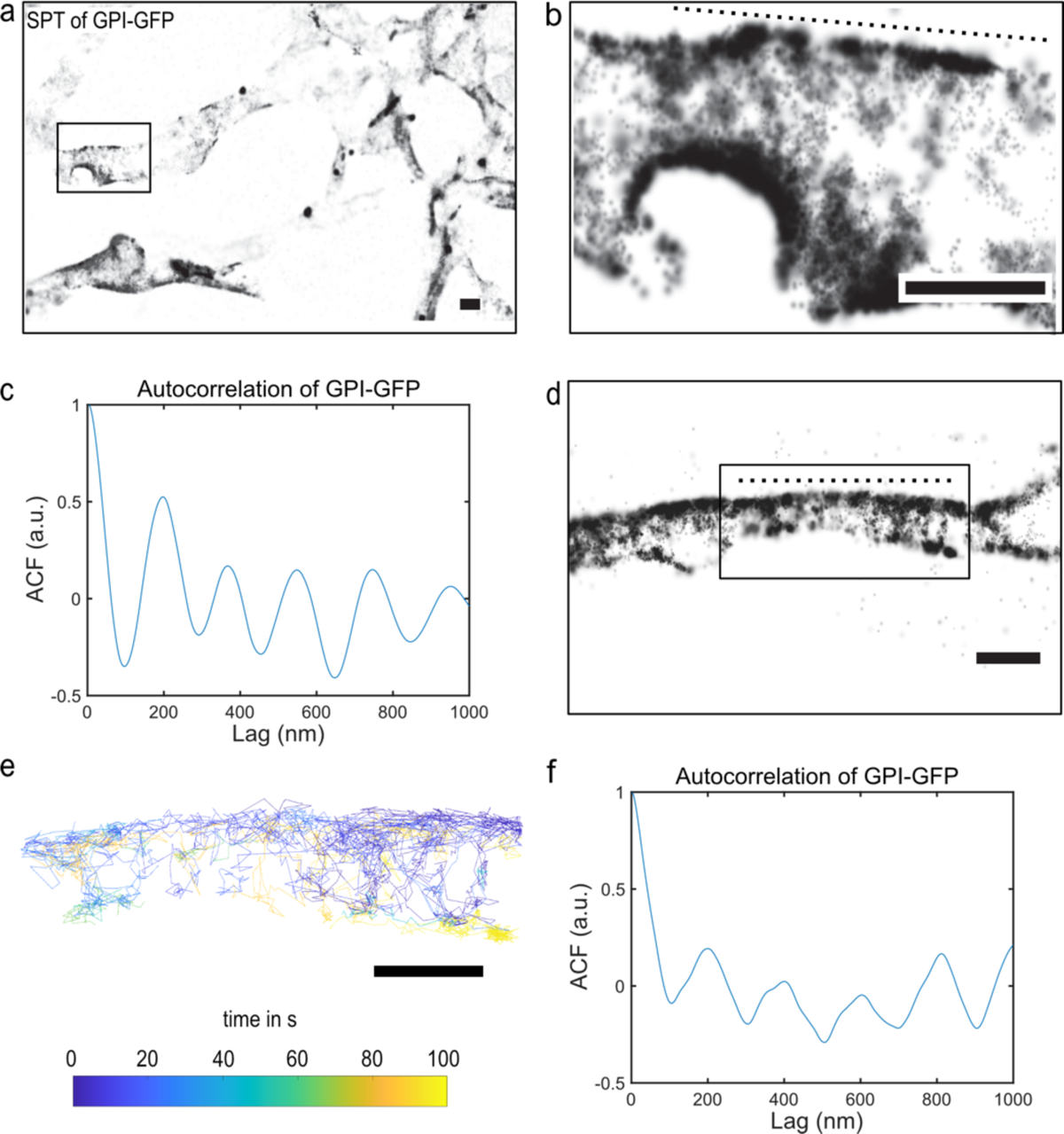
Membrane domains can be detected in progenitor-derived neuronal cells. **(a)** Reconstruction of SPT experiment (20000 frames, 200 Hz) of GPI-GFP tagged with QD’s in progenitor-derived neuronal cells. **(b)** Zoom in of box shown in (a) showing GPI-GFP forming membrane domains. **(c)** Autocorrelation along the dotted line along the neuronal process shown in (b) reveals periodicity of GPI-GFP domains at ∼200 nm. **(d)** Like (a) but different neuronal cell. **(e)** Single particle tracks color-coded for time of GPI-GFP motion of box shown in (d) reveals GPI-GFP membrane domains. **(f)** Like (c) but for neuron shown in (d). Scale bars are 500 nm.

Next, we investigated whether the membrane compartmentalization we detected for the peripheral outer leaflet membrane protein GPI-GFP could also be observed for the inner leaflet lipid-anchored Src family kinase Src-Halo (Boggon and Eck, 2004; Zhou et al., 2019) or the multi-spanning transmembrane cannabinoid receptor CB1 (Howlett, 2002; Zhou et al., 2019) coupled to YFP (Figure 4a). Indeed, after high-speed, high-density SPT, we could observe membrane compartmentalization for both molecules in the characteristic 200 nm spacing (Figure 4b, c), showing that also inner-leaflet and multi-spanning proteins are sensitive to the barriers located to the actin rings.

**Figure 4:**
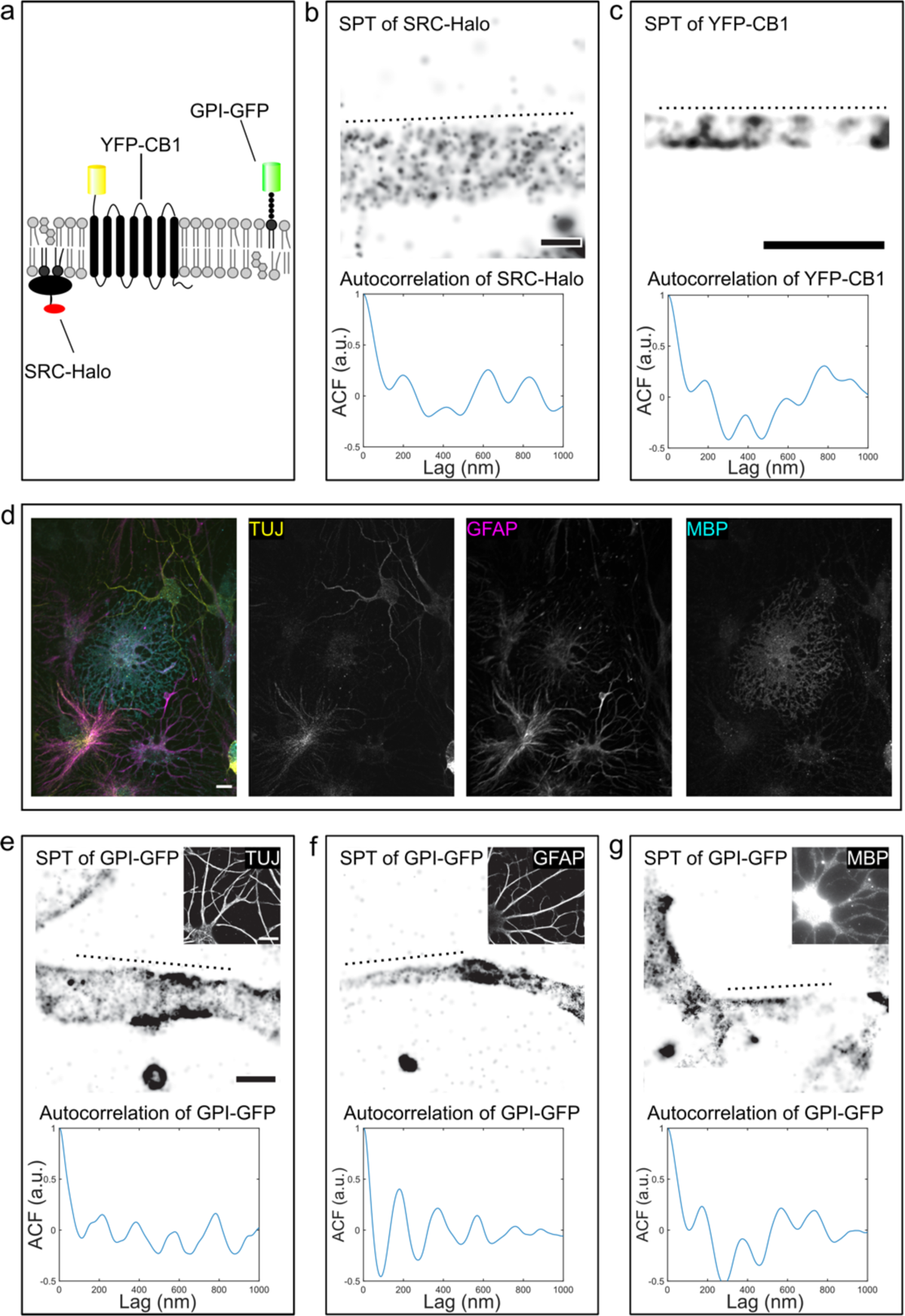
Membrane domains can be detected for different membrane molecule species and in several cell types of the neuronal lineage. **(a)** Schematic representation of molecule species that were investigated: the inner leaflet peripheral membrane protein: SRC-Halo, the transmembrane protein YFP-CB1 and the outer leaflet peripheral membrane protein GPI-GFP. **(b) (top)** Reconstruction of SPT experiment (20000 frames, 100 Hz) of SRC-Halo tagged JF635 **(bottom)** Autocorrelation along the dotted line along the neuronal process shown on top reveals periodicity of SRC-Halo domains at ∼200 nm. **(c) (top)** Reconstruction of SPT experiment (20000 frames, 200 Hz) of YFP-CB-1 tagged with QD’s in progenitor-derived neuronal cells **(bottom**) Autocorrelation along the dotted line along the neuronal process shown on top reveals periodicity of YFP-CB1 domains at ∼200 nm. **(d)** Confocal images of progenitor-derived neuronal cells stained for their lineage marker: **(left)** merge **(middle left)** neurons stained for TUJ **(middle right)** astrocytes stained for GFAP **(right)** oligodendrocytes stained for MBP. **(e) (top)** Reconstruction of SPT experiment (20000 frames, 200 Hz) of GPI-GFP tagged with QD’s in progenitor-derived neurons. Inset shows microscopy image of tracked cell stained for TUJ. Scale bar of inset is 10 µm. **(bottom)** Autocorrelation along the dotted line along the neuronal process shown on top reveals periodicity of GPI-GFP domains at ∼200 nm. **(f)** like (e) but for an astrocyte stained for GFAP (g) like (e) but for an oligodendrocyte stained for MBP. If not otherwise indicated, scalebars are 500 nm.

Sub-membrane actin rings spaced by spectrin tetramers have been described in a variety of cell types, including progenitor-derived astrocytes and oligodendrocytes (Hauser et al., 2018).

If our hypothesis was correct, these cells should thus also compartmentalize the motion of membrane proteins between actin rings. We thus differentiated progenitor cells into neuronal cells, astrocytes and oligodendrocytes (Figure 4d) and performed SPT experiments of GPI-GFP in all 3 cell types that were *a priory* identified via their morphology. After SPT experiments, cells were fixed and stained with markers of their respective lineage to confirm their cellular identity. When we then analyzed SPT localizations using autocorrelation analysis, we detected compartmentalization of GPI-GFP motion with a period of ∼200 nm for all cell types investigated here (Figure 4e-g). We concluded that membrane compartments are bounded by actin rings in many cell types.

We went on to perform correlated high-speed, high-density SPT of GPI-GFP and subsequent SMLM super-resolution imaging of actin in the same region of cells. When we overlayed the SPT and SMLM data from such experiments, we found that GPI-GFP molecules explored areas free from actin (Figure 5).

**Figure 5:**
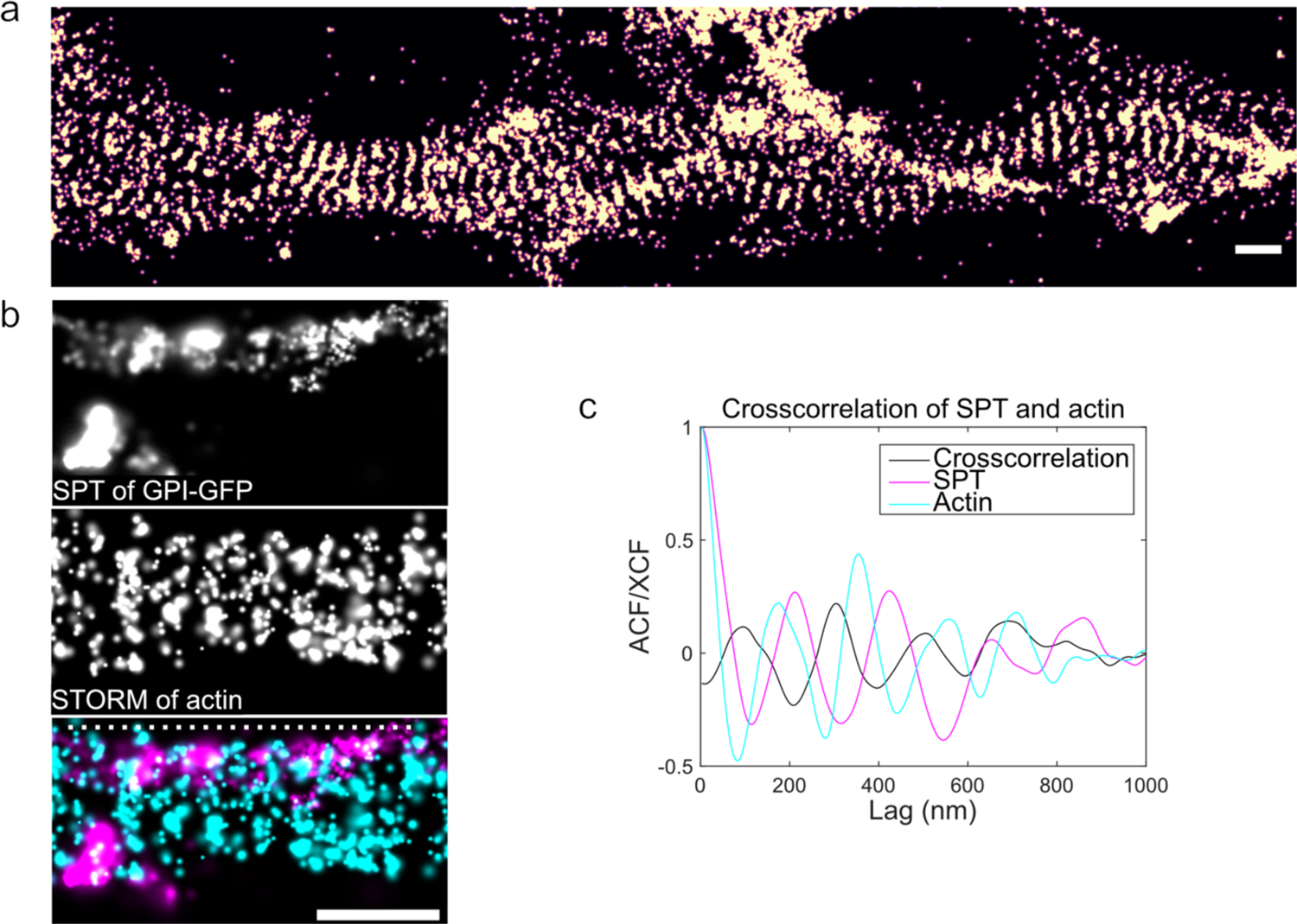
GPI-GFP accumulates between actin rings in progenitor-derived neuronal cells. **(a)** dSTORM reconstruction of actin in progenitor-derived neuronal cells stained with phalloidin. **(b)** Correlative SPT of GPI-GFP tagged with QD’s and dSTORM of actin experiments in progenitor-derived neuronal cells. **(top)** GPI-GFP was tracked using QD’s (5000 frames, 200 Hz). **(middle)** Cells were subsequently fixed and stained for actin using phalloidin. **(bottom)** Overlay of SPT and STORM reconstructions. GPI-GFP accumulates between actin rings. **(c)** Autocorrelation along dotted line in (b) reveals periodicity ∼200 nm of actin rings and GPI-GFP domains. Cross correlation of GPI-GFP domains and actin rings reveals that GPI-GFP accumulates between actin rings.

Finally, we decided to test, whether the actin rings, the location of membrane compartment boundaries, were indeed causing the membrane compartmentalization. To do so, we performed high-speed, high-density SPT experiments in live progenitor-derived neuronal cells before and after actin disruption by SwinholideA (SWIN A), a small molecule that cleaves actin filaments and inhibits actin polymerization (Spector et al., 1999; Klenchin et al., 2005; Vassilopoulos et al., 2020) (Figure S4). We reasoned that to detect compartmentalization in regions of neurons before treatment, but not after treatment, we would need an automated means of detecting such patterns to avoid bias due to insufficient sampling in our experiments or due to arbitrary choice of region of interest. To do so, we developed a software (Stripefinder) that would detect patterns in our simulated compartmentalized SPT data as in Figure 1g, but not in data generated from random walks (Figure S5a). With our software, we did not detect such patterns in accumulated high-speed, high-density SPT localizations in CV1 fibroblast cells that lack visible periodic actin structures (Figure S5b). Our software could however reliably detect regions in which autocorrelation analysis picked up stripes from accumulated high-speed, high-density SPT localizations (Figure S5c).

When we then analyzed accumulated high-speed, high-density SPT localizations from neuronal cells before and after treatment, we found many areas with periodic compartmentalization of GPI-GFP motion before treatment. We here specifically made sure that only areas that exhibited sufficient sampling density to make detection of rings possible in terms of localizations per area before and after drug addition were included in the analysis. However, in contrast, regions in which we detected stripes, did not exhibit them after SWIN A treatment (Figure 6 a-c). Similarly, we could not detect periodic localizations using autocorrelation analysis after SWIN A treatment when we reanalyzed areas in which stripes were detected before treatment (Figure 6d). We concluded that the disruption of actin rings by SWIN A reduces compartmentalization in the plasma membrane of progenitor-derived neuronal cells.

**Figure 6:**
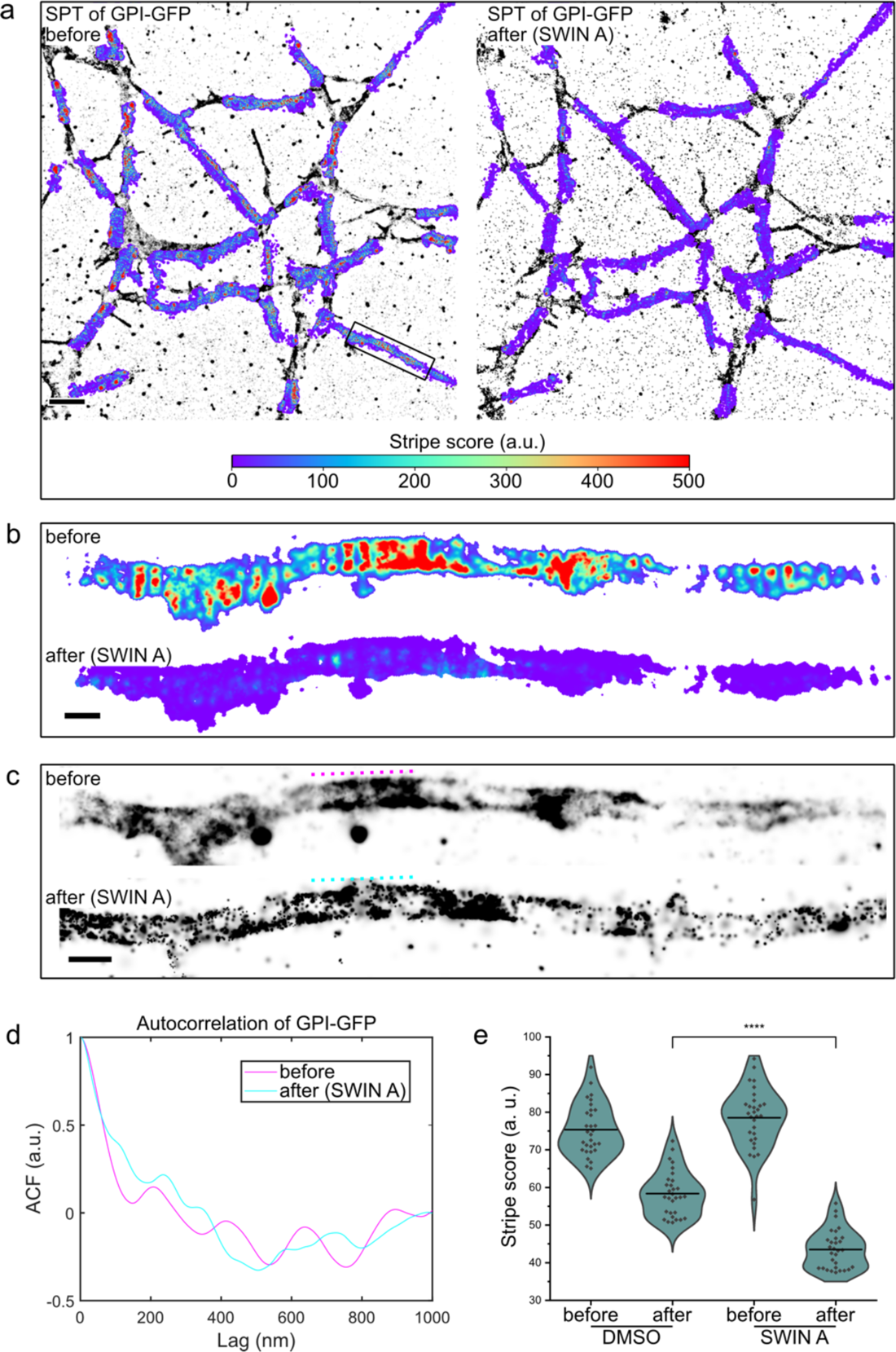
Actin disruption leads to loss of GPI-GFP accumulations. **(a)** Correlative SPT of GPI-GFP of progenitor-derived neuronal cells before and after SWIN A treatment. GPI-GFP was tracked using anti-GFP-NB-conjugated QD’s (5000 frames, 200 Hz). Cells were then treated with 1 µM SWIN A for 30 min. Afterwards, the same cell was tracked again. A local “stripe score” was then calculated based on the localizations via a flourier-filter based algorithm to detect periodicity at 200 nm. Shown are reconstructions of SPT experiments overlayed with heat maps of the local “stripe score”. After treatment, cells show a clear reduction in periodicity at 200 nm. Scalebar is 5 µm. **(b)** Heat map of stripe score of box in (a) before and after SWIN A treatment **(c)** Reconstructions of SPT experiments of box in (a) before and after SWIN A treatment **(d)** Autocorrelation plot of periodic area in (c) as indicated with dotted lines before and after SWIN A treatment. Periodicity ∼200 nm is lost after SWIN A treatment. **(e)** Experimental procedure as described in (a). SWIN A treated cells show a significantly reduced “stripe score” as compared to the control cells. Scalebars are 500 nm if not otherwise indicated.

## Discussion

We here used periodic actin rings in a variety of cell types as an experimental paradigm to test the hypothesis that sub-membrane actin structures can compartmentalize membrane protein motion. We find that actin rings compartmentalize the motion of transmembrane and peripheral membrane proteins linked to either leaflet of the plasma membrane by lipid anchors. This compartmentalization is independent of the presence of large clusters of immobilized membrane proteins between the actin rings such as found in the axon initial segment. We find that the actin rings are located a few nanometers below the plasma membrane and that their disruption leads to loss of compartmentalization. Taken together our results establish actin rings as a tractable experimental paradigm for the investigation of membrane compartmentalization.

We find that these actin rings, which are interconnected by spectrin tetramers in many neuronal cell types, are indeed closely apposed to the plasma membrane. Many transmembrane molecules are immobilized in the plasma membrane by cytoskeletal association and such sub-membrane actin rings, when spiked with many transmembrane domains may significantly obstruct motion of molecules across them ((Fujiwara et al., 2002) and Figure 1 and S1). Such an arrangement is the basis of the transmembrane “picket” model for membrane compartmentalization (Kusumi et al., 2005). This model has been proposed decades ago, however has been hard to prove experimentally, as cortical actin in live mammalian cells in very dynamic (Gowrishankar et al., 2012; Goswami et al., 2008) and in the presence of a thick actin cortex and stress fibers, membrane-apposed actin filaments are nearly impossible to detect (Clausen et al., 2017; Clark et al., 2013). However, we recently succeeded in pulling transmembrane proteins across the dorsal plasma membrane of cultured cells and found them to become stuck at the location of actin filamentous structures (Li et al., 2020). Indeed, it has long been observed that actin disruption influences membrane protein motion in cells (Andrade et al., 2015; Suzuki et al., 2005; Fujiwara et al., 2002) and lipids exhibit higher diffusion coefficients in blebs, in which no actin cortex is associated with the plasma membrane (Hiramoto-Yamaki et al., 2014).

Our work opens several important questions. We find that lipid-anchored molecules in the inner (Src-Halo) as well as the outer leaflet (GPI-GFP) become confined between actin rings, likewise transmembrane proteins (Cannabinoid receptor). Importantly, molecules remain mobile and unconfined inside compartments (Figure S3). Renner et al. have shown that tracking in 2D and under low acquisition speeds can lead to an underestimation of the observed diffusion coefficients due to the inherently 3-dimensional geometry of neurites. (Renner et al., 2011). We are however confident in the diffusion coefficients reported here, since all tracking data have been acquired under high acquisition speeds (5-10 ms) in 3D using a TIRF-microscope equipped with a biplane module.

Taken together, our observations support the transmembrane picket model. What could be the pickets serving as obstacles on actin rings? Recently, CD44 has been put forward to serve as picket to transmembrane diffusion in macrophages (Freeman et al., 2018), however, it is unclear if it is expressed in our cells. Recent mass spectrometric investigation of the periodic actin rings unfortunately did not produce novel actin ring-associated membrane proteins (Zhou et al., 2022). The only transmembrane protein reported to localize to actin rings is the potassium channel KV7.2 (D’Este et al., 2016), however it localizes to nodes of Ranvier and not ubiquitously over the axon or processes in neurons, let alone astrocytes or oligodendrocytes. In the future, it will be important to identify transmembrane proteins that may serves as pickets.

### What confines the molecules?

Actin disruption via SWIN A led to a reduction in membrane compartmentalization. This reduction was not complete, however the loss of periodic actin rings was neither, underlining again the exceptional stability of these structures. The actin rings remain exciting structures for further investigation. It was recently shown that they are formed from a particular, braided actin filament structure (Vassilopoulos et al., 2020) and that they are capable of changing their diameter to accommodate for large cargo (Wang et al., 2020), a capability that is likely mediated by non-muscle myosin II located to the rings (Berger et al., 2018; Zhou et al., 2022; Mikhaylova et al., 2020; Costa et al., 2020). Clearly, the actin rings influence membrane protein motion, but it remains to be shown, how such a ubiquitous method to compartmentalize membrane protein motion influences the biology of membrane proteins.

Since diffusion of membrane proteins is a principal biophysical property that strongly affects the kinetics of membrane protein reactions, and the rings are evolutionarily conserved at least in axons (He et al., 2016), their presence likely has significant functional consequences. It has been shown that receptor tyrosine kinase signaling is dependent on the periodic actin rings (Zhou et al., 2019), however, the mechanism behind this requirement remains unclear.

Likewise, neuronal adhesion molecules assume a periodic organization between actin rings in progenitor-derived oligodendrocytes (Hauser et al., 2018) and the rings are aligned between neighboring processes (Hauser et al., 2018; Zhou et al., 2022). The disruption of actin rings by spectrin knockdown leads to reduced axon bundling (Zhou et al., 2022), adhesion of dendrites to axons and even synapse formation, but it remains to be shown, if this is a specific effect or indeed the compartmentalization allows for more efficient lateral association of processes via transcellular adhesive protein interaction (Zhou et al., 2022).

Taken together, our work establishes the causal role of actin rings in membrane protein compartmentalization and offers a tractable experimental paradigm to investigate the control of membrane protein motion. In the future, new extremely high-speed single molecule tracking methods (Fujiwara et al., 2023) or MINFLUX microscopy (Balzarotti et al., 2017) will allow more insight into nanoscopic membrane protein compartmentalization.

## Material and methods

### Simulated SPT experiments

Simulated SPT experiments were generated using Fluosim (Lagardère et al., 2020). Model geometries where generated using custom Python scripts. Movies of 5 particles diffusing over the geometry were then generated using parameters similar to those observed in real data for either the actin model (crossing probability 30%, fluorophore spot size 150 nm, intensity 200, switch on/off rate 0.5 Hz, 5005 frames) or the channel model (crossing probability 0%, fluorophore spot size 150 nm, intensity 200, switch on/off rate 0.5 Hz, 5005 frames). Motion blur and Poisson noise were added to the movies using custom Python code. Picasso (Schnitzbauer et al., 2017) was then used to generate localizations and to render super-resolved reconstructions from the movies.

Additionally, to test the effect of different crossing probabilities on compartmentalization, super-resolved reconstructions were generated in Fluosim. The same parameters as described above were used for the actin model with varying crossing probabilities (0, 0.01, 0.1, 0.2 … 1). Ten reconstructions were generated per condition. The mean autocorrelation peak at 200 nm was then calculated from all reconstructions per condition using a custom MATLAB (MathWorks) script.

### Neuronal cell culture

Rat hippocampal neurons where cultured as described previously (Schmerl et al., 2022) in accordance with the Directive 2010/63/EU of the European Parliament on the protection of animals used for scientific purposes. Protocols for animal sacrifice were approved by the Regional Office for Health and Social Affairs in Berlin (“Landesamt für Gesundheit und Soziales; LaGeSo”) and the animal welfare committee of the Charité and carried out under permits T0280/10 and T-CH 0002/21. Rat hippocampal neurons were transfected on DIV1 with GPI-GFP using Lipofectamin 3000 (#L3000001, Thermofisher) following the manufacturer’s instructions.

Adult hippocampal neuronal progenitor cells (AHNPC) were cultured and differentiated into progenitor-derived neuronal cells as described previously (Hauser et al., 2018; Peltier et al., 2010). All experiments were performed on cells grown in a double layer in 35 mm x 10 mm plastic dishes (#627160, Cellstar) containing 25 mm coverslips (#631-1072, VWR). Both the plastic dishes and the glass coverslips were coated with ornithine/laminin. 300,000 AHNPCs were seeded onto the plastic dishes (#631-1072, VWR) and 200,000 cells on top of the coverslip before differentiation was induced.

Progenitor-derived neuronal cells were transfected on DIV1 with YFP-CB1 Lipofectamin 3000 (#L3000001, Thermofisher) following the manufacturer’s instructions.

### SPT

Lag16 anti-GFP-NB (Fridy et al., 2014) was labelled with biotin as described previously (Li et al., 2020). QD655-Streptavidin (#Q10121MP, Thermofisher) was incubated with biotinylated anti-GFP nanobodies in a 1:1 molar ratio for 10 min. Cells (DIV7 rat hippocampal neurons, DIV5-DIV14 progenitor-derived neuronal cells, CV-1) expressing either GPI-GFP or YFP-CB1 were then incubated with increasing amounts of QD-NB-conjugates (between 0.1 nM and 1 nM) in live cell imaging solution (#A14291DJ, Thermofisher) until desired QD density was reached depending on cell density and transfection efficiency. Cells were then washed with 1 mL live cell imaging solution.

Imaging was performed at a laser-power density of 0.6 kW*cm^-2^ using a 637 nm laser. Typically, 20000 frames were acquired per measurement at an exposure time of 5 ms. Progenitor-derived neuronal cells (DIV5-DIV14) expressing SRC-Halo were incubated with 0.25 nM JF635 in live cell imaging solution for 10 min. Afterwards, cells were washed with 1 mL live cell imaging solution. Imaging was performed at a laser-power density of 1.3 kW*cm^-2^ using a 637 nm laser. Typically, 20000 frames were acquired per measurement at an exposure time of 10 ms.

All samples were imaged using a Vutara 352 super-resolution microscope (Bruker) equipped with a Hamamatsu ORCA Flash4.0 sCMOS camera for super-resolution imaging and a 60× oil immersion TIRF objective with a numerical aperture of 1.49 (Olympus). Immersion Oil 23 C° (#1261, Cargille) was used.

Acquired raw data were localized using SRX (Bruker). Localizations were estimated by fitting single emitters to a 3D experimentally determined point spread function (PSF) under optimization of maximum likelihood. The maximum number of localization iterations performed before a given non-converging localization was discarded, was set to 40. PSFs were interpolated using the B-spline method. For reconstructions, localizations were rendered according to their radial precision. For single molecule tracking analysis, localizations were exported using SRX (Bruker) and tracked in 3D using the FIJI Mosaic tracker plugin (Linkrange: 3, Displacement: 2) (Sbalzarini and Koumoutsakos, 2005). Diffusion coefficients and alpha exponents were calculated in Mosaic. Tracks were filtered according to their alpha exponent (2>alpha>0). The mean alpha exponents and geometric mean diffusion coefficients of all tracks per measurement were calculated using a custom MATLAB (MathWorks) script.

### Correlative SPT / immunostainings

The sample holder containing samples of progenitor-derived neuronal cells (DIV5-DIV14) stably expressing GPI-GFP was fixated on the microscope using a 2-component adhesive (Picodent Twinsil, Picodent) using the manufacturer’s instructions to avoid sample drift during the staining process. Cells were tracked as described under section SPT and subsequently fixed on the microscope in 4% PFA (v/v)/PBS for 15 min. Then, samples were quenched in 50 mM NH_4_Cl/PBS for 30 min. Afterwards, permeabilization/blocking was performed with 1% BSA (w/v)/0.05 % Saponin (w/v)/4% Horse serum (v/v)/PBS for 45 min. Then, samples were incubated in a drop of Image-iT (#R37602, Thermofisher) for 30 min.

Samples were then stained with either rat anti-myelin basic protein (#ab7349, Abcam, 1:100), rabbit anti-GFAP (#ab7260, Abcam, 1:1,000) or mouse anti-Tuj (#SAB4700544, Sigma, 1:300) for 2 h in 1% BSA (w/v)/0.05 % Saponin (w/v)/4% Horse serum (v/v)/PBS. Afterwards, sample were washed 5 times with 1 mL 1% BSA (w/v)/0.05 % Saponin (w/v)/PBS. Then, samples where incubated with donkey anti-mouse AF647 (#A31571, Invitrogen. 1:500), goat anti-rabbit AF647 (#A21246, Invitrogen) or goat anti-rat AF647 (#A21247, Invitrogen, 1:500) for 1 h in 1% BSA (w/v)/0.05 % Saponin (w/v)/4% Horse serum (v/v)/PBS. Subsequently samples were washed thrice with 1 mL PBS and then the same cells were imaged there were tracked previously. Imaging was performed in PBS.

### Correlative SPT experiments under actin disruption

The sample holder containing coverslips of progenitor-derived neuronal cells (DIV5-DIV14) stably expressing GPI-GFP were fixated on the microscope using a (Picodent Twinsil, Picodent) using the manufacturer’s instructions to avoid sample drift during drug incubation time. Afterwards, cells were tracked as described under section SPT (n=30 from 3 independent experiments). Cells were then treated with either 1% DMSO or 1 µM SWIN A for 30 min. Subsequently, the same cells were tracked again without addition of new QD-NB-conjugates.

### Correlative SPT / STORM experiments

The sample holder containing samples of progenitor-derived neuronal (DIV5-DIV14) stably expressing GPI-GFP were fixated on the microscope using a (Picodent Twinsil, Picodent) using the manufacturer’s instructions to avoid sample drift during the staining process. Cells were tracked as described under section SPT. Afterwards, cells were fixed and stained for actin as described under section dSTORM. The cells that were previously tracked were then imaged for actin.

### Triple-color immunostainings

Samples of progenitor-derived neuronal cells (DIV5-DIV14) were fixed in 4% PFA (v/v)/PBS for 15 min. Then, samples were quenched in 50 mM NH_4_Cl/PBS for 30 min. Afterwards, permeabilization/blocking was performed with 1% BSA (w/v)/0.05 % Saponin (w/v)/4% Horse serum (v/v)/PBS for 45 min. Then, samples were incubated in drop of Image-iT (#R37602, Thermofisher) for 30 min. Samples were then stained with rat anti-myelin basic protein (#ab7349, Abcam, 1:100), rabbit anti-GFAP (#ab7260, Abcam, 1:1,000) and mouse anti-Tuj (#SAB4700544, Sigma, 1:300) overnight at 4°C in in 1% BSA (w/v)/0.05 % Saponin (w/v)/4% Horse serum (v/v)/PBS. Afterwards, samples were washed thrice with 1 mL 1% BSA (w/v)/0.05 % Saponin (w/v)/PBS for 5 min on shaker. Then, samples were incubated with donkey anti-mouse AF647 (#A31571, Invitrogen, 1:500), goat anti-rabbit AF568 (#A11036, Invitrogen) and goat anti-rat AF488 (#A11006, Invitrogen, 1:500) for 1 h in 1% BSA (w/v)/0.05 % Saponin (w/v)/4% Horse serum (v/v)/PBS. Subsequently samples were washed thrice with 1 mL PBS and imaging was performed in PBS on an inverted Olympus IX71 microscope equipped with a Yokogawa CSU-X1 spinning disk. A 60×/1.42 NA oil Olympus objective and a sCMOS camera (Hamamatsu) was used.

### AnkG immunostainings

DIV7 rat hippocampal neurons, DIV7 or DIV14 progenitor-derived neuronal cells were fixed, blocked and imaged as described under section triple-color immunostainings but were incubated with mouse anti-AnkG (#AB_10675130, Neuromab, 1:200) overnight at 4°C and subsequently with donkey anti-mouse AF647 (#A31571, Invitrogen, 1:800) for 1h.

### Generation of lentiviruses

Lentiviral transfer vectors containing the CDS for GPI-GFP or GFP-P2A-SRC-Halo and corresponding lentiviruses were generated at Viral Core Facility (Charité, Berlin).

### Generation of stable cell lines

For coating of dishes, culturing of cells and freezing of AHNPC’s see section neuronal cell culture. 1 mL of supernatant containing Lentivirus (GPI-GFP, or GFP-P2A-SRC-Halo) were added to 100,000 AHNPC’s growing on ornithine/laminin coated 6-well plates (#92406, TPP). Plates were centrifuged at 1000 rcf for 30 min. After 72 h of incubation cells were washed thrice with 1 mL PBS. Afterwards, cells were detached using 1 mL accutase for 20 min. Cells from 2 wells were resuspended in 10 mL PBS and centrifuged at 250 g for 5 min. Supernatant was discarded and cell pellet was resuspended in 10 mL DMEM/F-12 + N-2 and transferred to an ornithine/laminin-coated T75 dish (#90076, TPP). Cells were then extended and frozen.

### Cloning of LYN-Halo-pEGFP-N1

GPI-Halo-pEGFP-N1 was a gift from Akihiro Kusumi (OIST, Okinawa, Japan). pEGFP-N1 was a gift from Antony K Chen (Addgene plasmid # 172281; http://n2t.net/addgene:172281; RRID:Addgene_172281). A DNA fragment (BamHI-LYN-Halo-BsrGI) was generated by performing a PCR (forward primer: AAAAAGGATCCGCCACCATGGGCTGCATCAAGAGCAAGCGCAAGGACAACCTGAACGACGACGGCGTGGACACCGGTTTTTT, reverse primer: AAAAATGTACACTAGCCGGAAATCTCGAGCGTCG) using GPI-Halo-pEGFP-N1 as a template. The DNA fragment and the pEGFP-N1 backbone were digested using BsrGI-HF (#R3575SVIAL, NEB) and BamHI-HF (#R3136SVIAL, NEB) following the manufacturer’s instructions. The digested backbone and the digested fragment were then ligated and transformed into TOP10. Plasmid sequences were confirmed by Sanger sequencing.

### Cloning of SRC-Halo-pEGFP-N1

SRC-EGFP-pEGFP-N1 was a gift from Terence Dermody & Bernardo Mainou (Addgene plasmid # 110496; http://n2t.net/addgene:110496; RRID:Addgene_110496). A DNA fragment (BamHI-Src(c-term)-Halo(n-term) was generated by performing a PCR (forward primer: AAAGGATCCACCGGTCGCCACCATGGCAGAAATCG, reverse primer: AAAAATGTACACTAGC) using LYN-Halo-pGFPN1 as a template. The DNA fragment and the SRC-EGFP-pEGFP-N1 backbone were digested using BsrGI-HF (#R3575SVIAL, NEB) and BamHI-HF (#R3136SVIAL, NEB) following the manufacturer’s instructions. The digested backbone and the digested fragment were then ligated and transformed into TOP10. Plasmid sequences were confirmed by Sanger sequencing.

### Cloning of YFP-CB-1-pcDNA3

CB1-pcDNA3 was a gift from Mary Abood (Addgene plasmid # 13391; http://n2t.net/addgene:13391; RRID:Addgene_13391). L-YFP-GT46 was a gift from P. Keller (Max Planck Institute for Cell Biology and Genetics, Dresden, Germany).

The eYFP coding region containing a signal sequence was amplified from L-YFP-GT46 by performing a PCR using a primer pair of YFP_GT46_fwd (CAAGCTTGGTACCGAGCTCGGCCACCATGGAGCTCTTTTG) and YFP_GT46_rev (GGCCGTCTAAGATCGACTTCATGCCACTACTACTACTTCCACTACTACTACTTCCCTTGTACAGCTCGTCCATGCC). To assemble DNA fragment YFP_GT46_fwd-YFP_GT46_rev into CB-1-pcDNA3, primers were also designed to have 25-bp overlap at the junction regions. Afterward, CB-1-pcDNA3 was digested with BamHI-HF (#R3136SVIAL, NEB). To ligate the PCR product of YFP_GT46_fwd-YFP_GT46_rev into digested CB-1, Gibson Assembly Kit (#E5510S, NEB) was used following the manufacturer’s instruction. The resulting plasmid was transformed into TOP10. Plasmid sequences were confirmed by Sanger sequencing.

### STED microscopy

Progenitor-derived neuronal cells stably expressing GPI-GFP aged between DIV5 and DIV14 were stained live for actin using 1 µM SIR-actin (#SC001, Spirochrome) for 1 h at 37°C or stained for the plasma membrane using 10 nM Cholesterol-Starred (#STRED-0206, Abberior) for 10 min. For control experiment, cells were simultaneously incubated with Tetraspeck beads (1:500, #T7279, Invitrogen) for 1 h at 37°C. Cells were washed thrice with Live Cell Imaging Solution (#A14291DJ, Invitrogen). Then, samples were imaged in 1 mL Live Cell Imaging Solution (#A14291DJ, Invitrogen). STED microscopy was performed on an Abberior STED system with an 100×/1.4 NA oil uPlanSApo Olympus objective. STED imaging of SIR was performed with a 645 nm excitation and a 775 nm depletion pulsed laser. Detection was carried out with an avalanche photodiode (APD) with a 650 – 756 nm detection window. GFP was excited with a 485 nm laser and detected with an APD (498 to 551 nm). A single focal plane in the center of neuronal processes was imaged. Pixel size was set to 20 nm and pixel dwell time to 10 µs.

### Distance calculation between GPI-GFP and actin rings / plasma membrane

To measure the distance between actin rings and GPI-GFP (n=31 regions from 12 images) or between the plasma membrane (PM) and GPI-GFP (n=42 regions from 8 images) plot profiles along neuronal processes were taken in both the GPI-GFP channel (confocal) and the SIR/Starred channel (STED). The plot-profiles were analyzed using a custom MATLAB (MathWorks) script. In the plot profiles two peaks are apparent for the two sides of the PM / actin rings / GPI-GFP. The plot profiles where smoothed slightly by applying a moving mean. Afterwards the left and right peak in the plot profiles were identified by finding the maxima in the smoothed profiles. The diameter of the PM / actin rings / GPI-GFP was then calculated from the difference of the peaks. Half the difference of GPI-GFP and actin rings and half the difference of GPI-GFP and the PM was then used to calculate the corresponding distances. To control the accuracy of the distance calculation, the same analysis was performed on the distance of Tetraspeck (#T7279, Invitrogen) beads imaged in the green channel (confocal) and the far-red channel (STED) (n=47 regions from 7 images).

### dSTORM

Progenitor-derived neuronal cells (DIV5-DIV14) were fixed in 4% PFA / PBS for 15 min at 37°C. Then, samples were quenched in 50 mM NH4Cl/PBS for 30 min. Samples were subsequently stained with 2 µM Phalloidin-AF647 (#A22287, Thermofisher) in 1% BSA (w/v) / 0.05% saponine (v/v) / 4% horse serum (v/v) / PBS for 1 h. Afterwards, samples were washed 5 times with PBS. All samples were imaged using a Vutara 352 super-resolution microscope (Bruker, USA) equipped with a Hamamatsu ORCA Flash4.0 sCMOS camera for super-resolution imaging and a 60× oil immersion TIRF objective with a numerical aperture of 1.49 (Olympus, Japan). Immersion Oil 23 C° (#1261, Cargille, USA) was used. Samples were mounted onto the microscope in GLOX buffer (1.5% β-mercaptoethanol, 0.5% (v/w) glucose, 0.25 mg/ml glucose oxidase and 20 μg/ml catalase, 150 mM Tris-HCl pH 8.8). All imaging was performed at a laser-power density of 3.5 kW*cm^-2^ using a 637 nm laser. 10000 frames were acquired per measurement at an exposure time of 20 ms.

### Stripefinder software

We have developed a software pipeline for processing and segmentation of single molecule localization microscopy (SMLM) reconstructions and identifying periodic structures within. The pipeline has been implemented using Python and depends on NumPy (Harris et al., 2020), Pandas (McKinney, 2010) and SciPy (Virtanen et al., 2020) packages for computations and data analysis. Additionally, we have used the Scikit-Image (van der Walt et al., 2014) package for image processing operations.

Our designed pipeline is compatible with localizations from SMLM experiments in the form of sets of (x, y) coordinates with corresponding information on the point spread functions.

Matching similar regions: We implemented a convenient method for finding the required translations that result in the optimal match between two corresponding SMLM reconstructions (before SWINA treatment / after SWINA treatment). Reconstructions are loaded and rasterized into high-resolution bitmaps. This is followed with smoothing with identical Gaussian filters and segmentation using the Otsu’s thresholding method (van der Walt et al., 2014; N. Otsu, 1979). The resulting binary masks are superimposed and shifted in the x and y directions. Optimal shift in both directions is obtained via numerical minimization of the sum of absolute pixel-wise difference between the two images.

Segmentation of the region of interest: We have been interested in a functionality to automatically select features such as the edge regions of axons. Toward this goal, we start with the binary masks obtained from the previous step and transform them back into a sparse point-cloud. To reduce the number of points, we first find the skeletons of the binary masks via morphological operations. We sample the x and y coordinates of the remaining points and calculate their pairwise distance matrix. Nearest neighbors of each point are then found by applying a cut-off to the pairwise distances. We use the number of nearest neighbors of each point to assign a measure of connectivity to it. We consequently use the point-wise connectivity measure to construct a connectivity graph with highly-connected points as nodes. This allows us to segment the connectivity graph into ROI’s with a criterion that selects nodes belonging to an intact region, and relate the highlighted ROI’s back to the original image. We have used the regioncrops function of the Scikit-Image (van der Walt et al., 2014) package to uniquely label ROI’s in the final image.

We store the resulting information including coordinates of the point-clouds and the per-point ROI labels in a data structure for subsequent analysis. By iterating over the labels, we could select edges of axons, and (i) determine their orientation by measuring the slope of a simple least-squares linear fit, and (ii) apply appropriate scaling and cropping operations to focus on a region for further processing.

Identifying periodic structures via two-dimensional convolutions: In order to identify regions of the SMLM reconstructions that contain periodically occurring structures, we devised an approach based on two-dimensional convolution with a periodic kernel. The kernel comprises a series of elongated bivariate Gaussian functions that are positioned on a one-dimensional lattice, all oriented with a tilt angle θ with respect to the abscissa. Separation between the Gaussians is an input parameter of the method, and was chosen to mirror the expected separation between structures in the analyzed images.

Convolving this periodic kernel with the SMLM reconstructions results in local maxima for pixels around which the periodic tilted Gaussians best resembles the features in the image. To improve performance of the convolution operation, especially for large images, we implemented the convolution via Fourier transform. The well-known convolution theorem of the Fourier transform states that multiplication of signals in Fourier space corresponds to their convolution in real space. With the aid of fast Fourier transform (FFT) (Cooley and Tukey, 1965), the sequence of forward transform, pixel-wise multiplication with the kernel, and inverse transform has still better performance than the direct convolution in real space.

We processed the resulting convolved image further via a sweep with the discrete Laplace operator, which emphasizes the contrast in highlighted regions. By repeating the whole process for different values of the kernel tilt angle θ, we identified regions of the image containing periodic structures posed at different orientations.

## Author contributions

H.E. designed research. J.R., Ba.Se., P.S. performed research. S.B., J.R., M.S., F.N. developed analysis tools. J.R., S.B. analyzed data. F.B. provided training, equipment and reagents. Be.Sc, S.S. provided cell material. H.E., J.R. wrote the paper. All authors read and approved the final manuscript.

## Acknowledgements

This work was supported by Deutsche Forschungsgemeinschaft through SFB958/A04 and SFB1114/C03 and by the European Research Council through CoG 772230 “ScaleCell” and the Federal Ministry of Education and Research Germany through the grant “Deep-learning for XRM of organelle connectomics”. We thank all members of the Ewers laboratory for helpful discussions. Especially Purba Kashyap for input regarding data analysis and cloning and Jia Hui Li for input regarding data analysis and experimental procedures. We thank Thorsten Trimbuch and the team of the Viral Core Facility (Charité, Berlin) for the generation of the lentiviruses.

